## Supplementary Material for "IonBench: a benchmark of optimisation strategies for mathematical models of ion channel currents"

### A Problem Definitions

#### A.1 Problem data

All of the problems require generating synthetic data for the optimisation. For the Staircase and Loewe problems, these data are simulated current traces. These current traces are recorded at 2 kHz, while the Moreno $I_{Na}$ problem has varying frequencies to record its summary statistics. Additionally, the Staircase problems have i.i.d. $\sim \mathcal{N}(0, \sigma)$ noise added over the top of the current trace. The $\sigma$ for the noise is calculated as $\sigma = \frac{0.05}{n} \sum_{i=1}^{n} |I(t_i)|$, where $I(t_i)$ is the current at time step $t_i$.

#### A.2 Cost function

All of the problems use RMSE (Root Mean Squared Error) cost functions.

For the Moreno $I_{Na}$ problem, this is a weighted RMSE of the summary statistic curves. The error for each data point is weighted based on how many data points are in that curve. This means the final RMSE evenly weights errors in each of the summary statistics, rather than being biased towards the one with the most data points.

#### A.3 Parameters

The true parameters for each of the problems and the parameter bounds/sampling regions for the Staircase HH, MM, Loewe $I_{Kr}$, $I_{Kur}$ and Moreno $I_{Na}$ problems are given in Tables A.1, A.2, A.3, A.4, and A.5, respectively.

One thing to note for the Loewe problem rates, the specification of the model indicates the 'rates' only influence the timescale of the states, but not the steady state. The steady state is determined using separate, independent parameters.

| Index | True value | Bounds | Log-transformable |
|---|---|---|---|
| 1 | $2.26 \times 10^{-4}\,\mathrm{ms}^{-1}$ | $(10^{-7}, 10^{3})$ | Yes |
| 2 | $6.99 \times 10^{-2}\,\mathrm{mV}^{-1}$ | $(10^{-7}, 0.4)$ | No |
| 3 | $3.45 \times 10^{-5}\,\mathrm{ms}^{-1}$ | $(10^{-7}, 10^{3})$ | Yes |
| 4 | $5.462 \times 10^{-2}\,\mathrm{mV}^{-1}$ | $(10^{-7}, 0.4)$ | No |
| 5 | $8.73 \times 10^{-2}\,\mathrm{ms}^{-1}$ | $(10^{-7}, 10^{3})$ | Yes |
| 6 | $8.91 \times 10^{-3}\,\mathrm{mV}^{-1}$ | $(10^{-7}, 0.4)$ | No |
| 7 | $5.15 \times 10^{-3}\,\mathrm{ms}^{-1}$ | $(10^{-7}, 10^{3})$ | Yes |
| 8 | $3.158 \times 10^{-2}\,\mathrm{mV}^{-1}$ | $(10^{-7}, 0.4)$ | No |
| 9 | $0.1524\,\mu\mathrm{S}$ | $(0.02, 0.2)$ | No |

Table A.1: The parameters for the Staircase HH problem. The indices match those in Fig. 4. True values describe the parameters used to generate the data. Bounds are described as upper and lower bounds on parameters and define the parameter sampling region. The Staircase problems also ensure rate bounds are satisfied for sampled parameters. Log-transformable describes which parameters are transformed when log transforms are enabled.

#### A.4 Initial conditions

All problems are solved from steady state. This is solved both by `ionBench` and by `myokit`. The `ionBench` initial conditions are set up to allow automatic differentiation of the initial conditions with respect to the parameters, which is used to set the initial conditions of the sensitivity curves.

The implementation in `ionBench` does not include the same checks that are given in `myokit`, such as verifying the steady state is stable (which may be violated for some parameter combinations). This is why we also calculate the steady state using `myokit`. If there is any disagreement between `ionBench` and `myokit` in the steady state, then we instead simulate starting from a fully closed initial condition.

#### A.5 Solver tolerances

The ODE solver tolerances for each problem are given in Table A.6. The script used to verify the accuracy of the solver at these tolerances, and its output as a text file, are given in the GitHub repository.

| Index | True value | Bounds | Log-transformable |
|---|---|---|---|
| 1 | $2.0618 \times 10^{-1}\,\mathrm{ms}^{-1}$ | $(10^{-7}, 10^{3})$ | Yes |
| 2 | $1.12 \times 10^{-2}\,\mathrm{mV}^{-1}$ | $(10^{-7}, 0.4)$ | No |
| 3 | $4.209 \times 10^{-2}\,\mathrm{ms}^{-1}$ | $(10^{-7}, 10^{3})$ | Yes |
| 4 | $2.202 \times 10^{-2}\,\mathrm{ms}^{-1}$ | $(10^{-7}, 10^{3})$ | Yes |
| 5 | $3.65 \times 10^{-2}\,\mathrm{mV}^{-1}$ | $(10^{-7}, 0.4)$ | No |
| 6 | $4.1811 \times 10^{-1}\,\mathrm{ms}^{-1}$ | $(10^{-7}, 10^{3})$ | Yes |
| 7 | $2.23 \times 10^{-2}\,\mathrm{mV}^{-1}$ | $(10^{-7}, 0.4)$ | No |
| 8 | $1.3279 \times 10^{-1}\,\mathrm{ms}^{-1}$ | $(10^{-7}, 10^{3})$ | Yes |
| 9 | $6.03 \times 10^{-2}\,\mathrm{mV}^{-1}$ | $(10^{-7}, 0.4)$ | No |
| 10 | $8.094 \times 10^{-2}\,\mathrm{ms}^{-1}$ | $(10^{-7}, 10^{3})$ | Yes |
| 11 | $2.262 \times 10^{-4}\,\mathrm{ms}^{-1}$ | $(10^{-7}, 10^{3})$ | Yes |
| 12 | $3.99 \times 10^{-2}\,\mathrm{mV}^{-1}$ | $(10^{-7}, 0.4)$ | No |
| 13 | $4.15 \times 10^{-2}\,\mathrm{ms}^{-1}$ | $(10^{-7}, 10^{3})$ | Yes |
| 14 | $3.12 \times 10^{-2}\,\mathrm{mV}^{-1}$ | $(10^{-7}, 0.4)$ | No |
| 15 | $0.024\,\mu\mathrm{S}$ | $(0.02, 0.2)$ | No |

Table A.2: The parameters for the Staircase MM problem. The indices match those in Fig. 4. True values describe the parameters used to generate the data. Bounds are described as upper and lower bounds on parameters and define the parameter sampling region. The Staircase problems also ensure rate bounds are satisfied for sampled parameters. Log-transformable describes which parameters are transformed when log transforms are enabled.

| Index | True value | Bounds | Log-transformable |
|---|---|---|---|
| 1 | $3 \times 10^{-4}\,\mathrm{ms}^{-1}\,\mathrm{mV}^{-1}$ | $(3 \times 10^{-5}, 3 \times 10^{-3})$ | Yes |
| 2 | $14.1\,\mathrm{mV}$ | $(-45.9, 74.1)$ | No |
| 3 | $5\,\mathrm{mV}$ | $(0.5, 50)$ | Yes |
| 4 | $3.3328\,\mathrm{mV}$ | $(-56.6672, 63.3328)$ | No |
| 5 | $5.1237\,\mathrm{mV}$ | $(0.51237, 51.237)$ | Yes |
| 6 | 1 | $(0.1, 10)$ | Yes |
| 7 | $14.1\,\mathrm{mV}$ | $(-45.9, 74.1)$ | No |
| 8 | $6.5\,\mathrm{mV}$ | $(0.65, 65)$ | Yes |
| 9 | $15\,\mathrm{mV}$ | $(-45, 75)$ | No |
| 10 | $22.4\,\mathrm{mV}$ | $(2.24, 224)$ | Yes |
| 11 | $2.9411765 \times 10^{-2}\,\mathrm{nS}\,\mathrm{pF}^{-1}$ | $(2.9411765 \times 10^{-3}, 2.9411765 \times 10^{-1})$ | Yes |
| 12 | $138.994\,\mathrm{mmol}$ | $(13.8994, 1389.94)$ | Yes |

Table A.3: The parameters for the Loewe $I_{Kr}$ problem. The indices match those in Fig. 4. True values describe the parameters used to generate the data. Bounds are described as upper and lower bounds on parameters and define the parameter sampling region. If the parameters are log-transformable, then they are sampled from a log-uniform distribution rather than a uniform distribution. Log-transformable describes which parameters are transformed when log transforms are enabled. Units differ from Loewe et al. [1] to ensure model consistency.

| Index | True value | Bounds | Log-transformable |
|---|---|---|---|
| 1 | $0.65\,\mathrm{ms}^{-1}$ | $(0.065, 6.5)$ | Yes |
| 2 | $10\,\mathrm{mV}$ | $(-50, 70)$ | No |
| 3 | $8.5\,\mathrm{mV}$ | $(0.85, 85)$ | Yes |
| 4 | $30\,\mathrm{mV}$ | $(-30, 90)$ | No |
| 5 | $59\,\mathrm{mV}$ | $(5.9, 590)$ | Yes |
| 6 | $2.5$ | $(-57.5, 62.5)$ | No |
| 7 | $82\,\mathrm{mV}$ | $(22, 142)$ | No |
| 8 | $17\,\mathrm{mV}$ | $(1.7, 170)$ | Yes |
| 9 | $30.3\,\mathrm{mV}$ | $(-29.7, 90.3)$ | No |
| 10 | $9.6\,\mathrm{mV}$ | $(0.96, 960)$ | Yes |
| 11 | $3$ | $(0.3, 30)$ | Yes |
| 12 | $1\,\mathrm{ms}^{-1}$ | $(0.1, 10)$ | Yes |
| 13 | $21$ | $(-39, 81)$ | No |
| 14 | $185\,\mathrm{mV}$ | $(125, 245)$ | No |
| 15 | $28\,\mathrm{mV}$ | $(2.8, 280)$ | Yes |
| 16 | $158\,\mathrm{mV}$ | $(98, 218)$ | No |
| 17 | $16\,\mathrm{mV}$ | $(1.6, 160)$ | Yes* |
| 18 | $99.45\,\mathrm{mV}$ | $(39.45, 159.45)$ | No |
| 19 | $27.48\,\mathrm{mV}$ | $(2.748, 274.8)$ | Yes |
| 20 | $3$ | $(0.3, 30)$ | Yes |
| 21 | $0.005\,\mathrm{nS\,pF}^{-1}$ | $(-59.995, 60.005)$ | No |
| 22 | $0.05\,\mathrm{nS\,pF}^{-1}$ | $(0.005, 0.5)$ | Yes |
| 23 | $15\,\mathrm{mV}$ | $(-45, 75)$ | No |
| 24 | $13\,\mathrm{mV}$ | $(1.3, 130)$ | Yes |
| 25 | $138.994\,\mathrm{mmol}$ | $(13.8994, 1389.94)$ | Yes |

Table A.4: The parameters for the Loewe $I_{Kur}$ problem. The indices match those in Fig. 4. True values describe the parameters used to generate the data. Bounds are described as upper and lower bounds on parameters and define the parameter sampling region. If the parameters are log-transformable, then they are sampled from a log-uniform distribution rather than a uniform distribution. Log-transformable describes which parameters are transformed when log transforms are enabled. Units differ from Loewe et al. [1] to ensure model consistency. * This parameter was originally labelled as additive [1], but its appearance in the model and the originally used parameter bounds suggest it should be treated as multiplicative, so we allow it to be log transformed.

| Index | True value | Bounds | Log-transformable |
|---|---|---|---|
| 1 | $7.6178 \times 10^{-3}\,\mathrm{ms}$ | $(5.71 \times 10^{-3}, 9.52 \times 10^{-3})$ | Yes |
| 2 | $32.764\,\mathrm{mV}$ | $(24.6, 41)$ | No |
| 3 | $5.8871 \times 10^{-1}$ | $(4.42 \times 10^{-1}, 7.36 \times 10^{-1})$ | Yes |
| 4 | $1.5422 \times 10^{-1}$ | $(1.16 \times 10^{-1}, 1.93 \times 10^{-1})$ | Yes |
| 5 | $2.5898\,\mathrm{ms}$ | $(1.94, 3.24)$ | Yes |
| 6 | $8.5072\,\mathrm{mV}$ | $(6.38, 10.6)$ | No |
| 7 | $1.3760 \times 10^{-3}$ | $(1.03 \times 10^{-3}, 1.72 \times 10^{-3})$ | Yes |
| 8 | $2.888$ | $(2.17, 3.61)$ | Yes |
| 9 | $3.2459 \times 10^{-5}\,\mathrm{ms}^{-1}$ | $(2.43 \times 10^{-5}, 4.06 \times 10^{-5})$ | Yes |
| 10 | $9.5951\,\mathrm{mV}$ | $(7.20, 12)$ | No |
| 11 | $1.3771\,\mathrm{ms}^{-1}$ | $(1.03, 1.72)$ | Yes |
| 12 | $21.126\,\mathrm{mV}$ | $(15.8, 26.4)$ | No |
| 13 | $11.086\,\mathrm{ms}^{-1}$ | $(8.31, 13.9)$ | Yes |
| 14 | $43.725\,\mathrm{mV}$ | $(32.8, 54.7)$ | No |
| 15 | $4.1476 \times 10^{-2}$ | $(3.11 \times 10^{-2}, 5.18 \times 10^{-2})$ | Yes |
| 16 | $2.0802 \times 10^{-2}$ | $(1.56 \times 10^{-2}, 2.60 \times 10^{-2})$ | Yes |

Table A.5: The parameters for the Moreno $I_{Na}$ problem. The indices match those in Fig. 4. True values describe the parameters used to generate the data. Bounds are described as upper and lower bounds on parameters (rounded to 3 significant figures here) and define the parameter sampling region. Log-transformable describes which parameters are transformed when log transforms are enabled.

The solver tolerances are set such that three conditions are satisfied. The first two conditions concern the magnitude of the solver noise (calculated as the standard deviation of the cost around a parameter vector) across the parameter space, and the third concerns the cost threshold.

The first condition samples parameters across the whole parameter space, calculates the local standard deviation in the cost for small ($\mathcal{O}(10^{-12})$) perturbations in parameters, and verifies the median is below $10^{-7}$. The second condition is that the $75^{\text{th}}$ percentile of the standard deviation of the cost is below $10^{-6}$. The third condition verifies that points near to the true parameters always satisfy the cost threshold for the problem.

Note that only the Staircase problems use the ODE solver for the cost function evaluation. The other problems use an analytical solver, and use the specified tolerances only for solves with sensitivities, as analytical solves with sensitivities are not available in `myokit`.

| Problem | Abs. tolerance | Rel. tolerance |
|---|---|---|
| Staircase HH | $10^{-5}$ | $10^{-5}$ |
| Staircase MM | $10^{-6}$ | $10^{-6}$ |
| Loewe $I_{Kr}$ | $10^{-6}$ | $10^{-6}$ |
| Loewe $I_{Kur}$ | $10^{-8}$ | $10^{-8}$ |
| Moreno $I_{Na}$ | $10^{-7}$ | $10^{-7}$ |

Table A.6: Bootstrapped significance. The absolute and relative solver tolerances used for each problem.

### B Model solve time

One of the assumptions in the results is that the model always takes the same amount of time to solve, regardless of the approach. It is possible that some approaches or optimisers propose more or less stiff parameters which may lead to a corresponding change in the time to solve the model for those parameters.

In addition to recording the number of function evaluations in `ionBench`, we also record the time to solve the model. These are reported in Figs B.1 and B.2 for the solves without and with sensitivities, respectively. It is clear that the average solve time for the models is independent of the choice of approach, meaning it is reasonable to use function evaluations in place of model solve time for the ERT calculations.

There is an outlier in the Moreno $I_{Na}$ problem cost timings in Fig. B.1. This point corresponds to the Sachse2003b approach. During all the optimisations with this approach, the first two points were correctly solved, the third point resulted in a failed solved (giving a NaN cost), and the remaining solves were all attempted at NaN parameters. All optimisations resulted in 48 model solves (all without sensitivities), but because these solves failed, they were able to fail quickly. Since the average model solve times for each problem are weighted by the number of solves for each approach, this outlier has not affected the average model solve time for the Moreno $I_{Na}$ problem significantly.

### C Significance

#### C.1 Bootstrapping algorithm

In this section, we describe the bootstrapping algorithm used to verify the significance of the results in `ionBench`.

Algorithm 1 describes how a single bootstrapped ERT sample is generated. Algorithm 2 describes how we compare the ERT samples for the best approach against each other approach.

#### C.2 Significance

All approach-problem pairs that were not significantly worse than the Wilhelms2012b approach paired with that problem were rerun with an unbounded number of maximum runs. They (both the approach to be evaluated and Wilhelms2012b) were run against 50 parameter samples (10 for the Staircase HH problem) and the significance was evaluated. If the Wilhelms2012b approach was not found to be significantly better or significantly worse, then a further 50 (or 10) parameters were sampled and evaluated against. This continued until the Wilhelms2012b approach was significantly better or worse than each other approach-problem pairing.

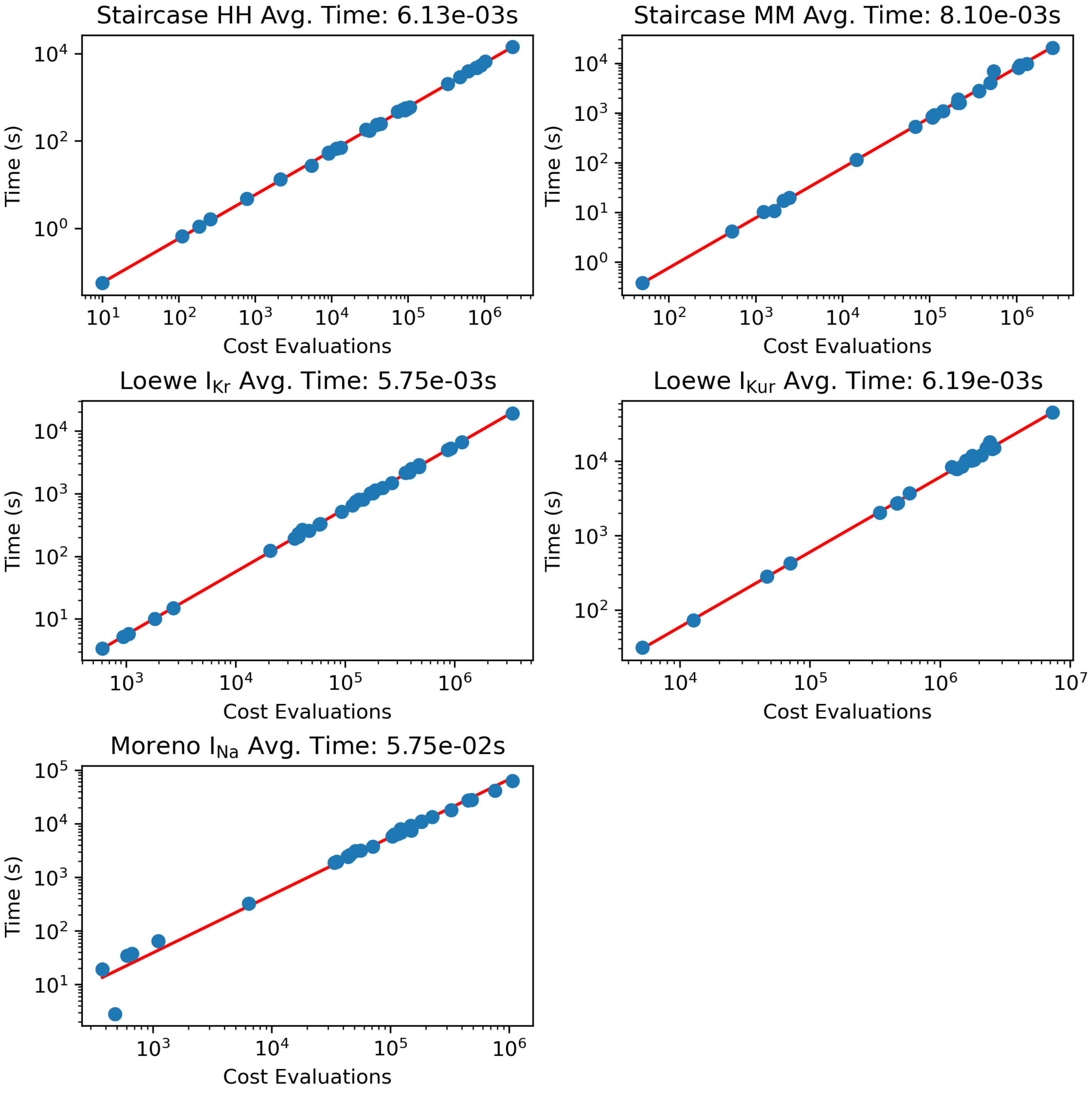


Figure B.1: Plots of the total time spent solving the model (without sensitivities) against the number of function evaluations for each approach (cumulative across the $n_{\text{Runs}}$).

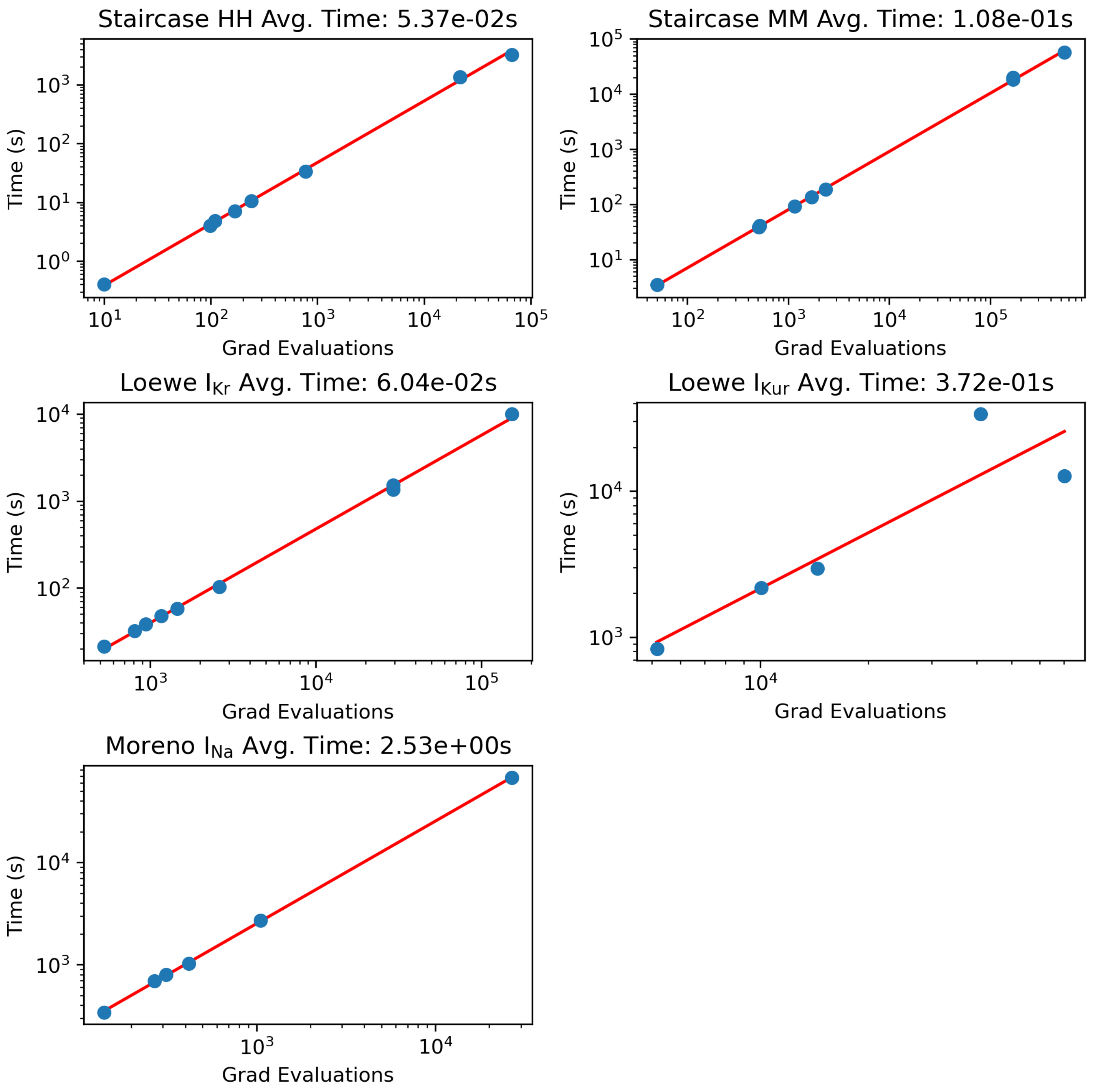


Figure B.2: Plots of the total time spent solving the model (with sensitivities) against the number of function evaluations for each approach (cumulative across the $n_{\text{Runs}}$).

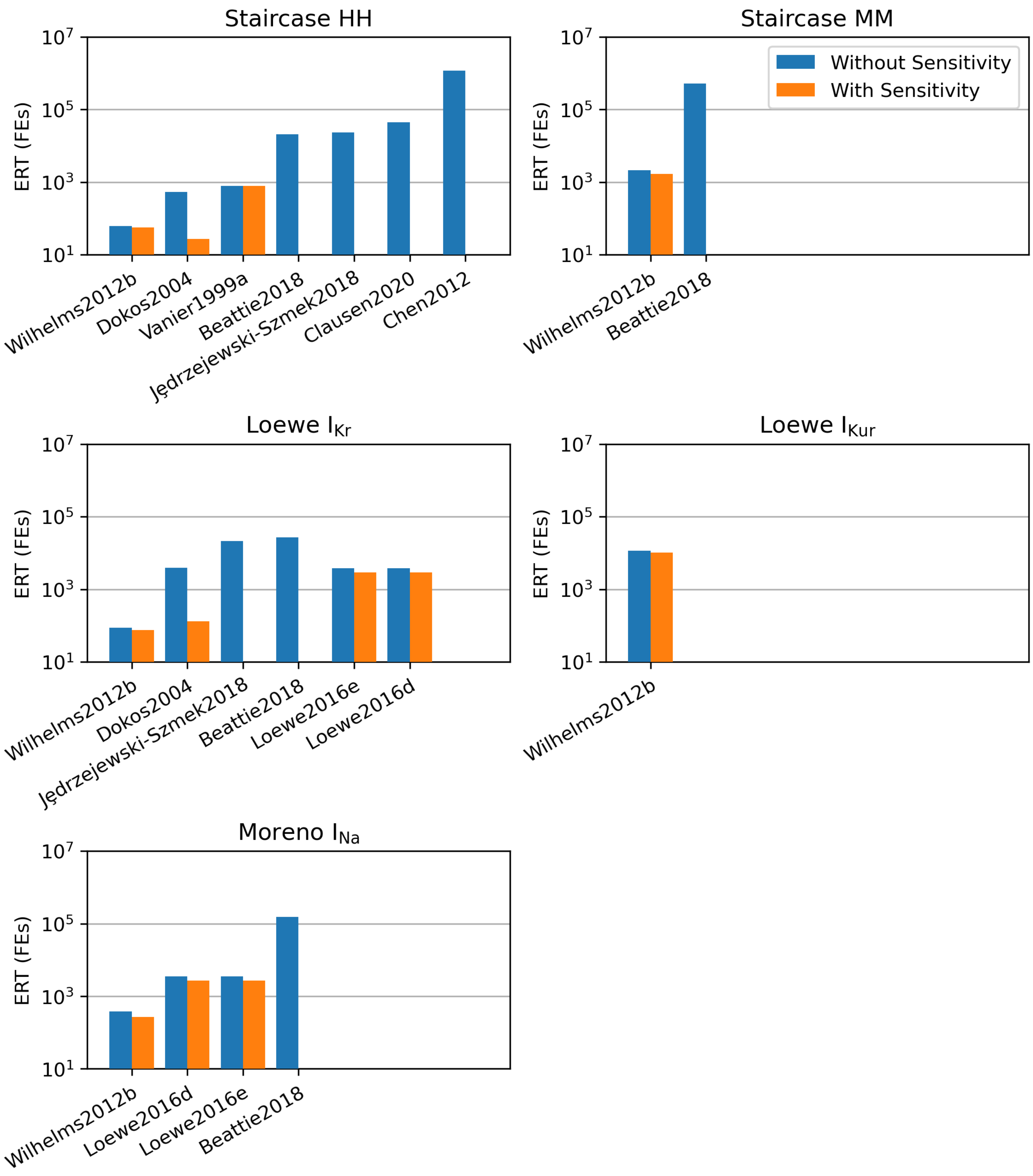


Figure B.3: Plots of ERT, separated into solves with and without sensitivities. The total ERT, as shown in Fig. 6, is given as the without sensitivity ERT plus the with sensitivity ERT after being scaled by the time ratio in Table 2.

**Algorithm 1:** Draw a single sample of ERT using data from a particular approach and problem

**Input:** t: List of times for an approach. s: List of success/failures for an approach
**Output:** ERT: Sampled ERT for bootstrapping
$nRuns = \text{len}(t)$;
`Draw bootstrap sample of optimisation runs`
$mask = \text{choose}(from = 1 : nRuns, count = nRuns, replacement = True)$;
$s_{bs} = s[mask]$;
$t_{bs} = t[mask]$;
`Sample bootstrapped success rate`
$p \sim Beta(\alpha = 0.5 + \text{sum}(s_{bs} == True), \beta = 0.5 + \text{sum}(s_{bs} == False))$;
$p_{bs} = \text{drawSample}(p)$;
`Calculate average success and fail times`
**if** $\text{sum}(s_{bs} == success) > 0$ **then**
  $T_s = \text{mean}(t_{bs}[s_{bs} == success])$;
**else**
  `Choosing a single value maximises the bootstrapped variances when we are most unsure about` $T_s$
  $T_s = \text{choose}(from = t_{bs}, count = 1)$;
**end**
**if** $\text{sum}(s_{bs} == fail) > 0$ **then**
  $T_f = \text{mean}(t_{bs}[s_{bs} == fail])$;
**else**
  `Choosing a single value maximises the bootstrapped variances when we are most unsure about` $T_f$
  $T_f = \text{choose}(from = t_{bs}, count = 1)$;
**end**
$ERT = T_s + T_f \times (1 - p_{bs})/p_{bs}$;

Table C.1 presents the number of runs required for each approach before the results became significant. In all cases, except for the Dokos2004-Staircase HH pairing, the results are significant in favour of the Wilhelms2012b approach.

The parameters used for the significance testing for a given problem are consistent across approaches, but are different from the initial $n_{\text{Run}}$ parameters.

There are also a couple of edge cases, where slightly different methods were used to improve computation times. In one case (Chen2012 approach for the Staircase MM problem), the batch size was unsuitable due to long computational times, so a smaller batch size of 5 was used. While generally, the approaches were tested against the same parameters as Wilhelms2012b, for the Groenendaal2015 approach (and Staircase MM problem), 2500 parameters were used for Wilhelms2012b (to reduce its uncertainty) and Groenendaal2015 was then run in batches (of size 10) until significance. Otherwise, Groenendaal2015 likely would not have seen significance until around 400 evaluated parameters. Similarly, 5000 parameters were used for Wilhelms2012b and Bueno-Orovio2008 was allowed to continue until significance (in this case, over 5000 parameters).

### D Profile Likelihood

Profile likelihood plots for the problems are given in Figs D.1, D.2, D.3, D.4, and D.5. Since not all of the problems have noise, we report the RMSE cost rather than the likelihood (negative log likelihood is linearly related to the square of the cost), though still referred to here as profile likelihood curves.

These figures are generated through ionBench and use the TRR optimiser. The code to generate the profile likelihood plots is given in the GitHub repository.

Since the Staircase MM shows very flat profile likelihood curves from some parameters, a figure where each plot is placed on its own scale is given in Fig. D.6. This clearly highlights a shift in the global minimum. This shift is the result of the noise added to the data, where the model can attempt to reproduce some small bias by moving to parameters offset from those that generated the data. This has been previously reported [2], albeit with a smaller bias in parameters (likely due to using a stiffer model/parameterisation here). We have verified resampling the noise moves the bias in the minimum, with the distribution centred on the data-generating parameters, and shrinking the noise reduces the bias in the parameters (although to slowly

**Algorithm 2:** The bootstrapping algorithm for comparing against the best approach.

**Input:** B: number of bootstrapped samples per approach; T[i]: list of run times for approach i; S[i]: list of boolean success/failures for approach i.

**Output:** sig: List of length $n = len(T[i]) = len(S[i])$ boolean indicating if approach 1 is significantly better than approach $i$.

```
Sort T and S by ERT such that T[1] and S[1] correspond to the approach that gives the lowest ERT;
Get number of approaches
n = len(T);
Generate bootstrapped samples for each approach
for i in 1 to n do
    Sample B ERTs for approach i
    for b in 1 to B do
        ERT[i][b] = Algorithm 1(T[i], S[i]);
    end
end
Compare approaches for significance
for i in 1 to n do
    if i==1 then
        No point in comparing approach 1 against itself
        sig[i] = N/A;
    else
        count = 0;
        for b_1, b_2 in 1 to B do
            if ERT[i][b_1] < ERT[1][b_2] then
                count = count + 1;
            end
        end
        count = count/B^2;
        if count > 0.05 then
            Approach i is better in over 5% of samples
            sig[i] = False;
        else
            Approach 1 is better in over 95% of samples
            sig[i] = True;
        end
    end
end
```

Table C.1: **Number of runs required for significance.**

| Approach | Staircase HH | Staircase MM | Loewe $I_{Kur}$ | Moreno $I_{Na}$ |
|---|---|---|---|---|
| Balser1990a | 40 | 1200 | N/A | 50 |
| Balser1990b | A/S | 400 | A/S | 50 |
| Vanier1999a | A/S | 600 | 750 | 50 |
| Vanier1999b | A/S | 400 | A/S | 50 |
| Vanier1999c | A/S | 3750 | 300 | 50 |
| Vanier1999d | A/S | 400 | 350 | 50 |
| Sachse2003a | A/S | 600 | A/S | N/A |
| Sachse2003b | A/S | 400 | 450 | 350 |
| Dokos2004 | 80 | 2350 | A/S | 150 |
| Bueno-Orovio2008 | 160 | 7950 | 450 | 50 |
| Seemann2009a | A/S | 400 | A/S | A/S |
| Liu2011 | A/S | 400 | A/S | A/S |
| Chen2012 | A/S | N/A | N/A | 45 |
| Davies2012 | A/S | 400 | A/S | 50 |
| Groenendaal2015 | A/S | 20 | A/S | A/S |
| Loewe2016d | A/S | A/S | N/A | 100 |
| Loewe2016e | A/S | A/S | N/A | 100 |
| Moreno2016 | A/S | 400 | A/S | 50 |
| Beattie2018 | A/S | 600 | A/S | 50 |

The number of runs required to show significance. Duplicate approaches (see Table 1), approaches that were already significantly worse for all problems, or problems where Wilhelms2012b was already always significantly better are not shown. A/S means that the Wilhelms2012b approach was already significantly better and further runs were not needed. N/A means the approach-problem paring failed to complete the initial $n_{\mathrm{Run}}$ optimisations in the 1 week limit, or otherwise failed (e.g. the Carins2017 approach is incompatible with negative parameters). Cells highlight in blue show where an approach was significantly better than Wilhelms2012b, otherwise the Wilhelms2012b approach was significantly better.

to allow both noise and minimal bias in the parameter minimum).

The cost thresholds are calculated from the profile likelihood curves of each problem. We begin by identifying the cost of the profile likelihood curves at ±5% around the true parameters (data-generating). We then look at taking the minimum of these values, across all profile likelihood curves for a given problem, with some exceptions. We want to ignore any unidentifiable parameters, and also ensure the cost threshold is far enough away from the minimum that approaches that implement a function tolerance termination criteria do not abort early (which would require editing hyperparameters to resolve).

The unidentifiable parameters that are removed are parameters 1, 2, 4, 6, 7, 11, 12, 13, 14, 16, and 20 of the Loewe $I_{Kur}$ problem (implemented as ignoring parameters if their perturbed cost is below $10^{-14}$). The limit of $10^{-14}$ is specific to the profile likelihood curves here and will not work in general for other unknown problems.

We also need to ignore any parameters where the perturbed cost (at the ±5% positions) is below the cost of the data-generating parameters. This is relevant for the Staircase MM problem, which sees a significant noise-induced bias in the global minimum. If this step was not done, then the cost threshold could end up arbitrarily close to the global minimum, depending on the cost of the perturbed parameters.

Next, we find the minimum of the perturbed costs, across all profile likelihood curves for each problem, ignoring any points that meet the above criteria.

Finally, we want to ensure there is some buffer room in the cost threshold. To do this, we will round the cost threshold up to 3 significant figures. If this step was not taken, then some problems, particularly Staircase MM where the cost function is very flat, would trigger many different termination criteria (function and gradient tolerances) too early, and these approaches would never be able to identify a minimum without changing hyperparameters, even if they reproduce the data almost perfectly.

The Loewe $I_{Kur}$ problem sees many unidentifiable parameters. Of the 25 profile likelihood curves in Fig. D.4, 11 show unidentifiability up to optimisation tolerances.

The Moreno $I_{Na}$ problem, shown in Fig. D.5, shows some optimisation difficulties in generating the profile likelihood plots. In all cases, either the profile likelihood curves at the ±5% points that are used to determine the cost thresholds appear to be accurate, or a small portion of the smooth profile likelihood curve can be seen and is well above the cost threshold. This suggests that these difficulties (which we believe are due to the post processing to calculate the summary statistics) are unlikely to influence the cost threshold calculation.

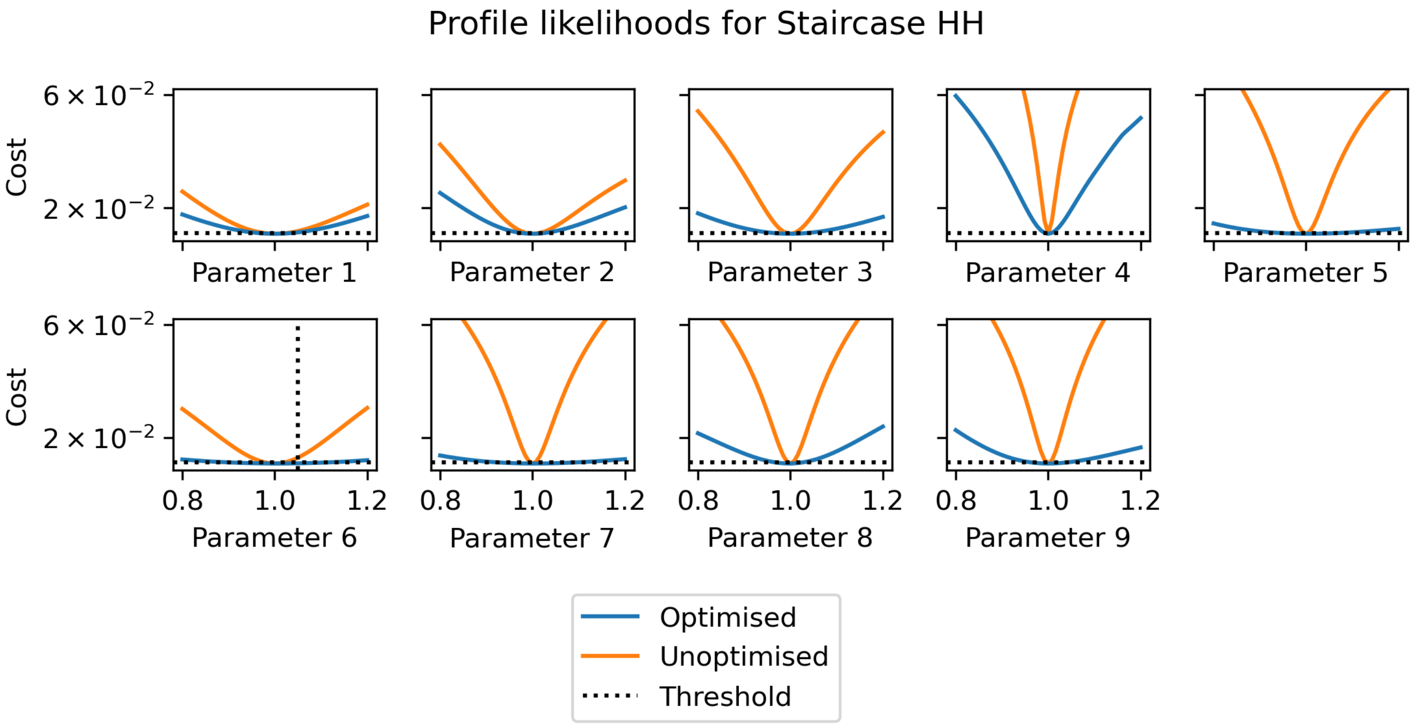


Figure D.1: Profile likelihood plots of the Staircase HH problem. The RMSE cost is given rather than likelihood, so the maximum likelihood estimates are given at the minimum cost. The true parameters are at $x = 1$. Both the profile likelihood curve (blue) and an unoptimised cost surface slice (orange) are shown. The cost threshold is shown as a horizontal dotted line, and the point which determines its value is labelled with a vertical dotted line.

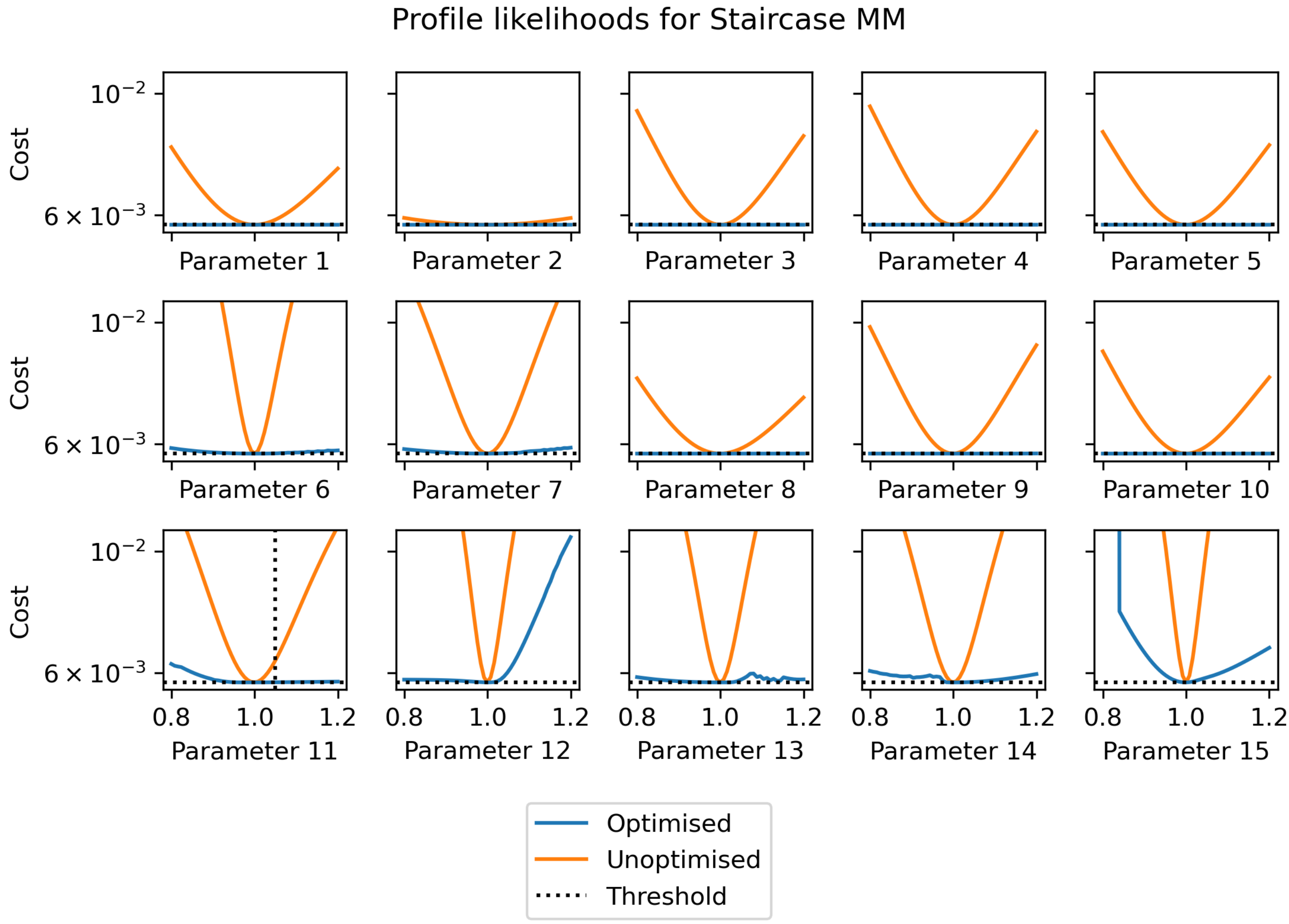


Figure D.2: Profile likelihood plots of the Staircase MM problem. The RMSE cost is given rather than likelihood, so the maximum likelihood estimates are given at the minimum cost. The true parameters are at $x = 1$. Both the profile likelihood curve (blue) and an unoptimised cost surface slice (orange) are shown. The cost threshold is shown as a horizontal dotted line, and the point which determines its value is labelled with a vertical dotted line.

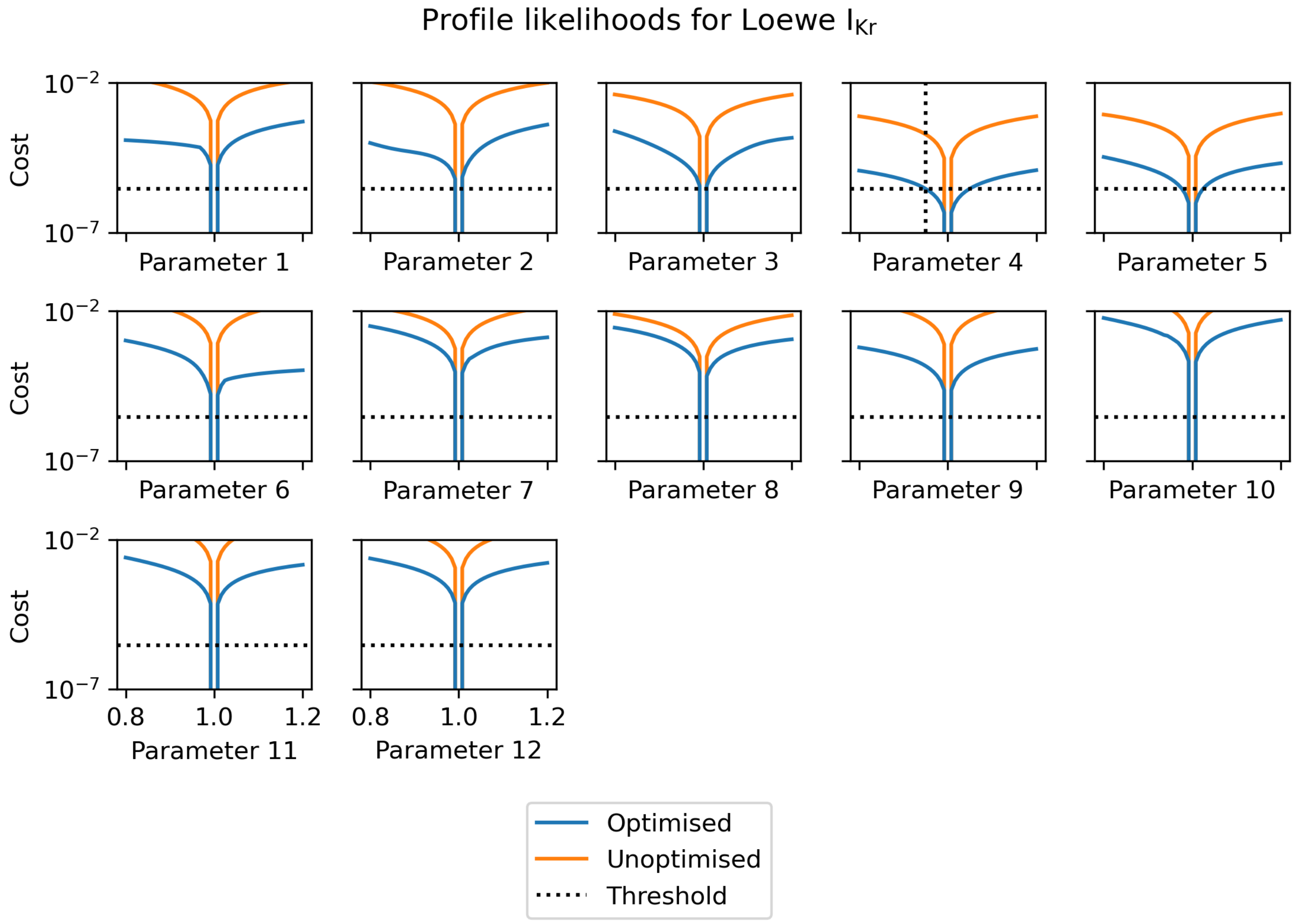


Figure D.3: Profile likelihood plots of the Loewe $I_{Kr}$ problem. The RMSE cost is given rather than likelihood, so the maximum likelihood estimates are given at the minimum cost. The true parameters are at $x = 1$. Both the profile likelihood curve (blue) and an unoptimised cost surface slice (orange) are shown. The cost threshold is shown as a horizontal dotted line, and the point which determines its value is labelled with a vertical dotted line.

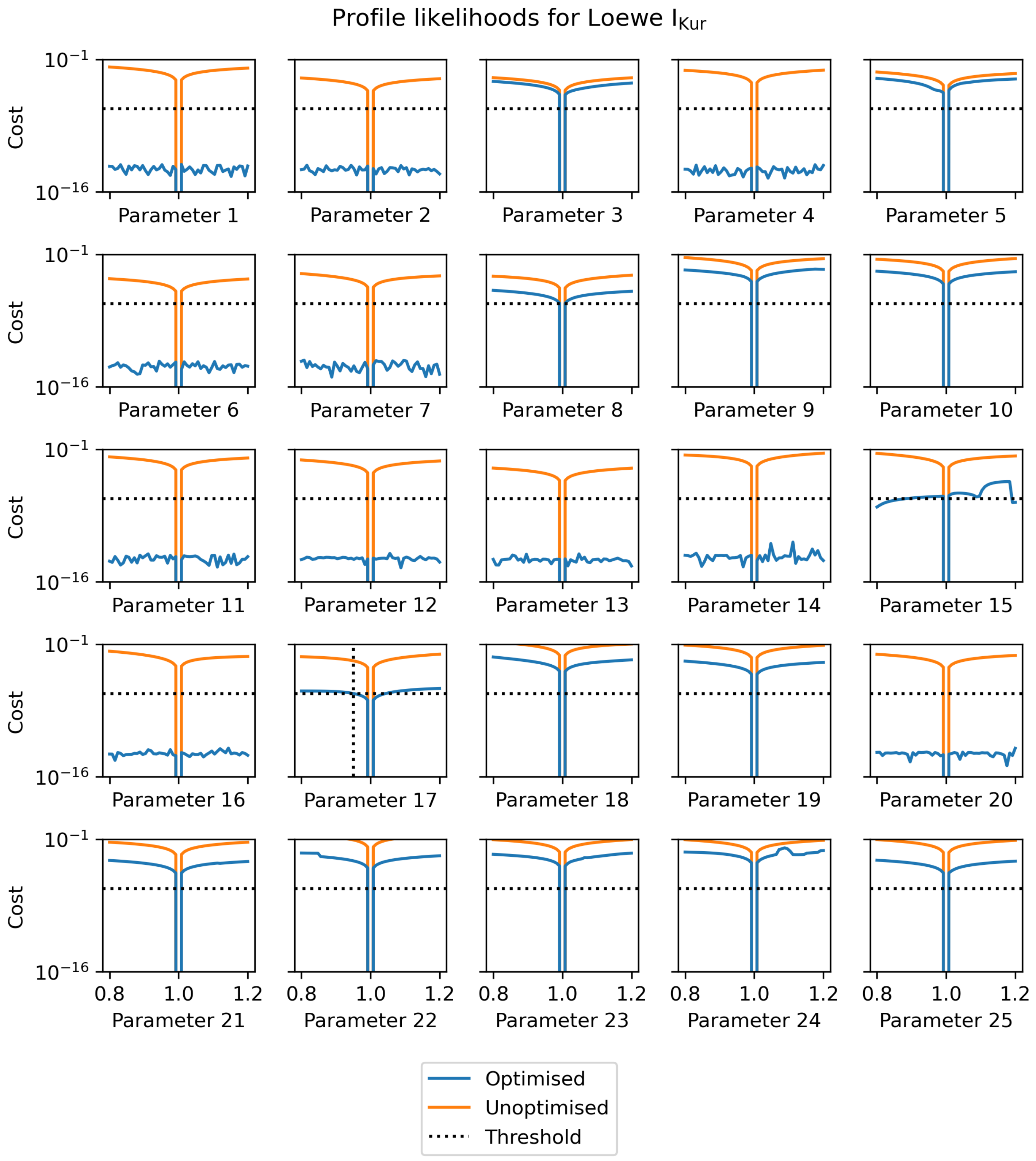


Figure D.4: Profile likelihood plots of the Loewe $I_{Kur}$ problem. The RMSE cost is given rather than likelihood, so the maximum likelihood estimates are given at the minimum cost. The true parameters are at $x = 1$. Both the profile likelihood curve (blue) and an unoptimised cost surface slice (orange) are shown. The cost threshold is shown as a horizontal dotted line, and the point which determines its value is labelled with a vertical dotted line.

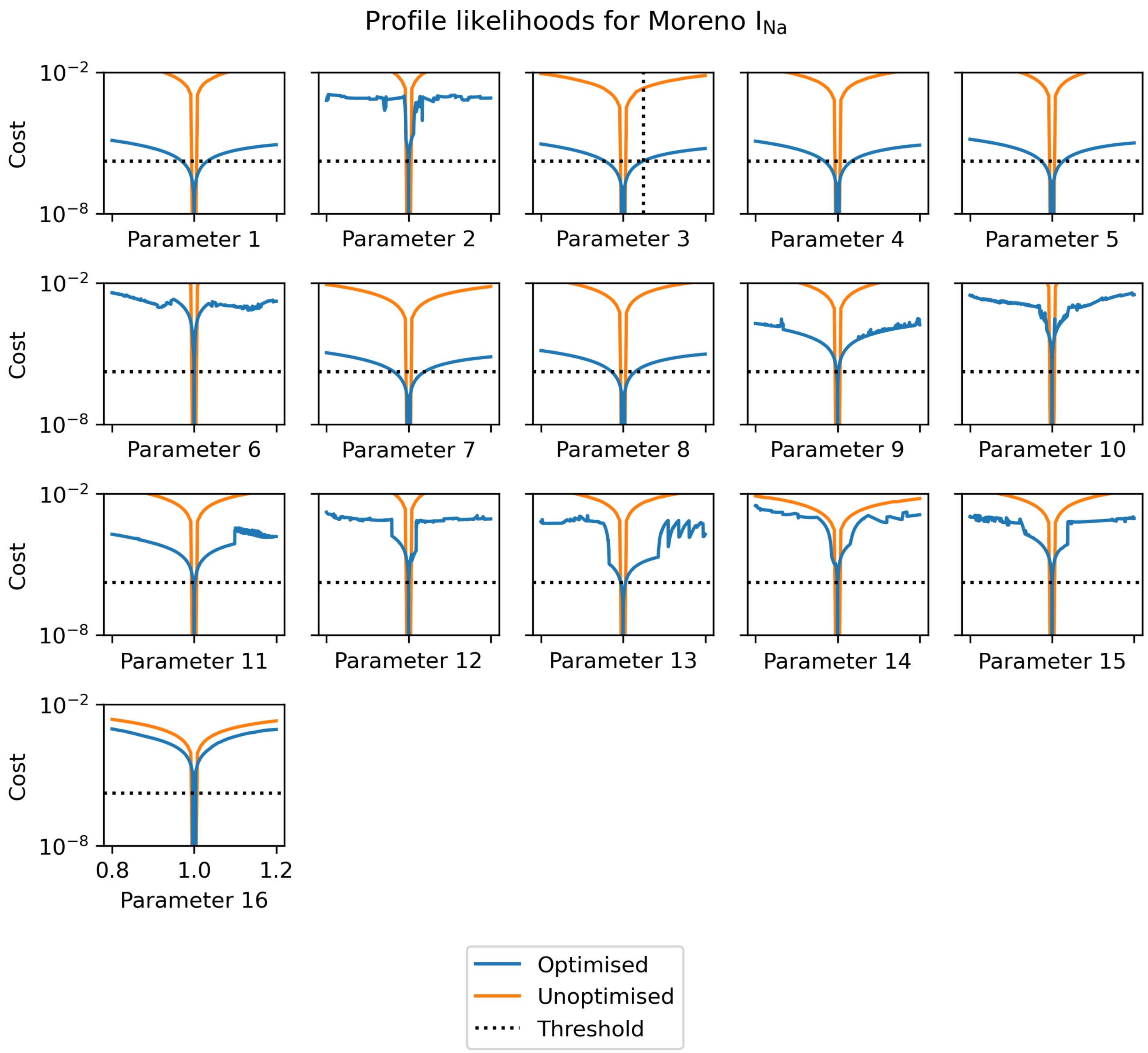


Figure D.5: Profile likelihood plots of the Moreno $I_{Na}$ problem. The RMSE cost is given rather than likelihood, so the maximum likelihood estimates are given at the minimum cost. The true parameters are at $x = 1$. Both the profile likelihood curve (blue) and an unoptimised cost surface slice (orange) are shown. The cost threshold is shown as a horizontal dotted line, and the point which determines its value is labelled with a vertical dotted line.

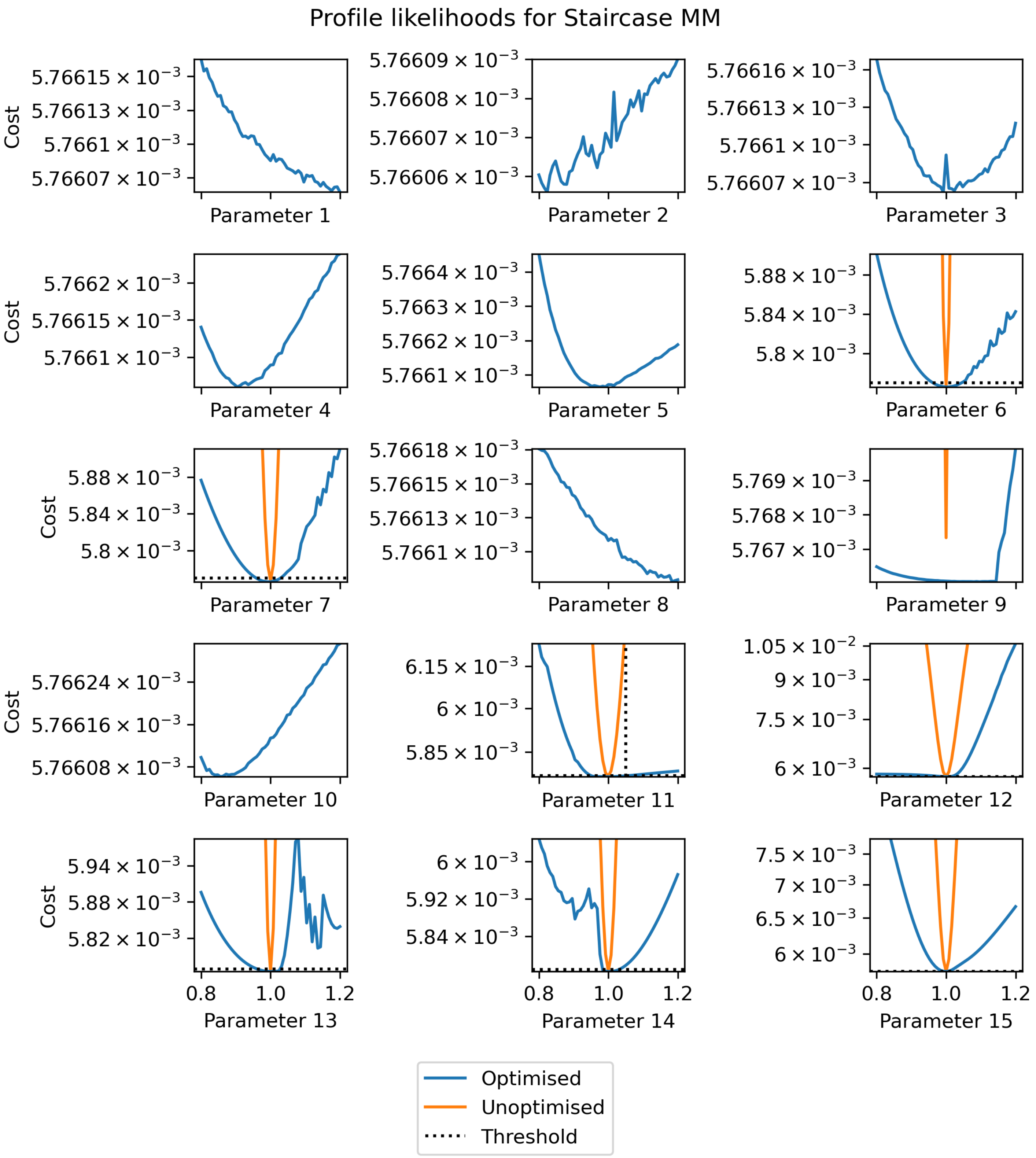


Figure D.6: Duplicate of Fig. D.2, but each subplot is on an independent scale. Profile likelihood plots of the Staircase MM problem. The RMSE cost is given rather than likelihood, so the maximum likelihood estimates are given at the minimum cost. The true parameters are at $x = 1$. Both the profile likelihood curve (blue) and an unoptimised cost surface slice (orange) are shown. The cost threshold is shown as a horizontal dotted line, and the point which determines its value is labelled with a vertical dotted line.
